# Biochemical characterization of actin assembly mechanisms with ALS-associated profilin variants

**DOI:** 10.1101/2022.01.05.475096

**Authors:** Xinbei Liu, Morgan L. Pimm, Brian Haarer, Andrew T. Brawner, Jessica L. Henty-Ridilla

## Abstract

Eight separate mutations in the actin-binding protein profilin-1 have been identified as a rare cause of amyotrophic lateral sclerosis (ALS). Profilin is essential for many neuronal cell processes through its regulation of lipids, nuclear signals, and cytoskeletal dynamics, including actin filament assembly. Direct interactions between profilin and actin monomers inhibit actin filament polymerization. In contrast, profilin can also stimulate polymerization by simultaneously binding actin monomers and proline-rich tracts found in other proteins. Whether the ALS-associated mutations in profilin compromise these actin assembly functions is unclear. We performed a quantitative biochemical comparison of the direct and formin-mediated impact for the eight ALS-associated profilin variants on actin assembly using classic protein-binding and single-filament microscopy assays. We determined that the binding constant of each profilin for actin monomers generally correlates with the actin nucleation strength associated with each ALS-related profilin. In the presence of formin, the A20T, R136W, Q139L, and C71G variants failed to activate the elongation phase of actin assembly. This diverse range of formin-activities is not fully explained through profilin-PLP interactions, as all ALS-associated variants bind a formin-derived PLP peptide with similar affinities. However, chemical denaturation experiments suggest that the folding stability of these profilins impact some of these effects on actin assembly. Thus, changes in profilin protein stability and alterations in actin filament polymerization may both contribute to the profilin-mediated actin disruptions in ALS.

## Introduction

Amyotrophic lateral sclerosis (ALS) is a fatal neurodegenerative disease characterized by progressive decline and loss of motor neuron function (Cleveland and Rothstein, 2001). There is no effective therapeutic or cure. No detailed mechanism explains ALS onset or motor neuron degeneration, though impairments to RNA metabolism and protein homeostasis are commonly explored (Ling et al., 2013; Van Damme et al., 2017). The role of the neuronal cytoskeleton in ALS is understudied despite its essential roles in biological processes linked to disease onset, particularly: DNA/RNA repair, protein degradation, neuronal excitability, and vesicular transport (Ajroud-Driss and Siddique, 2015; Castellanos-Montiel et al., 2020; Ling et al., 2013). Several specific mutations in five genes that directly regulate components of the neuronal cytoskeleton have been identified as causative in sporadic and familial ALS patients, including the microtubule motor protein kinesin-5a, neurofilaments, dynactin-1, tubulin (i.e., the building blocks of microtubules), and profilin-1 (Castellanos-Montiel et al., 2020; Gregory et al., 2020). With the exception of profilin, these candidates strongly suggest the involvement of intermediate filaments and microtubules as ALS progresses. While profilin regulates microtubule dynamics in ALS, dogma suggests that ALS-linked mutations in profilin are the mechanistic link between disruptions in actin dynamics and ALS onset (Boopathy et al., 2015; Figley et al., 2014; Giampetruzzi et al., 2019; Henty-Ridilla et al., 2017; Schmidt et al., 2021; Wu et al., 2012)

Profilin is a critical regulator of neuronal growth, development, and dendritic spine formation (Da Silva et al., 2003; Lambrechts et al., 2006; Lee et al., 2013; Michaelsen-Preusse et al., 2016; Neuhoff et al., 2005; Witke et al., 1998). It is regulated by diverse lipids and participates in numerous signaling pathways that regulate proteostasis, RNA transcription, nuclear export, and microtubule dynamics (Moens and Coumans, 2015; Pimm et al., 2020; Pinto-Costa and Sousa, 2019). Profilin is best characterized as an actin monomer binding protein that suppresses the spontaneous nucleation of actin filaments (Krishnan and Moens, 2009; Pimm et al., 2020). In contrast, profilin also stimulates actin assembly by simultaneously binding to actin monomers and the poly-*L*-proline (PLP) rich sequences of other proteins like formins or Ena/VASP (Henty-Ridilla and Goode, 2015; Rotty et al., 2015; Suarez et al., 2015; Suarez and Kovar, 2016). Through these combined functions, profilin effectively regulates most known mechanisms of cellular actin assembly (Funk et al., 2019; Skruber et al., 2020, 2018; Suarez and Kovar, 2016).

ALS-associated mutations in profilin are commonly thought to disrupt actin filament assembly mechanisms, either through direct interactions with actin monomers or indirectly through their association with actin assembly promoting proteins like Ena/VASP and formin (Alkam et al., 2017; Giampetruzzi et al., 2019; Schmidt et al., 2021). In a synthetic lethality screen from yeast, the C71G, M114T, and G118V ALS-related variants fail to complement the loss of wild-type profilin (Figley et al., 2014). Mammalian cell immunoprecipitation experiments indicate these variants directly bind to actin monomers and specific formin proteins (i.e., mDia1, mDia2, and FMNL1) (Figley et al., 2014; Giampetruzzi et al., 2019; Schmidt et al., 2021). Several studies suggest that at least a subset of these ALS-linked profilin proteins mildly destabilize actin dynamics in cells (Boopathy et al., 2015; Figley et al., 2014; Freischmidt et al., 2015; Schmidt et al., 2021; Wu et al., 2012). Other analyses imply that C71G, T109M, M114T, G118V, and to a lesser extent E117G misfold in ALS-pathology (Boopathy et al., 2015; Del Poggetto et al., 2016; Figley et al., 2014; Kiaei et al., 2018; Pereira et al., 2019; Sadr et al., 2021). Whether the ALS-related profilin variants directly disrupt actin binding or formin-based assembly mechanisms has not been fully explored.

Given the increasing evidence of an active role for profilin in the regulation of cytoskeletal proteins beyond actin assembly (i.e., tubulin dimers and microtubules), we sought to more completely characterize the ALS profilin variants with respect to their actin-based interactions (Pimm et al., 2021; Pimm and Henty-Ridilla, 2021). This would allow us to determine which profilin variants are (and which are not) significantly altered for activities with different cytoskeletal systems. To achieve this goal, we purified wild-type profilin, the eight ALS-associated profilin variants (A20T, C71G, T109M, M114T, E117G, G118V, R136W, Q139L), and profilin mutants with impaired actin binding (R88E and H120E) or deficient poly-*L*-proline binding (Y6D) (Ezezika et al., 2009; Kovar et al., 2006; Lu and Pollard, 2001; Suetsugu et al., 1998). Using fluorescence polarization, we determine the binding constants (k_D_) for each protein and actin monomers. We assess the effects of each profilin variant for defects in actin filament assembly mechanisms using bulk fluorescence assays and through direct visualization of single actin filaments in super resolution microscopy assays. We investigate the effects of folding stability in regulating actin assembly in chemical denaturation/refolding assays. Finally, we explore the ability of each variant to stimulate formin-based actin polymerization and determine the binding affinities for each protein for a PLP-peptide derived from the formin mDia1. We find that the ALS-profilins vary widely in their ability to interact with actin and to stimulate formin-based actin polymerization, suggesting that profilin-based ALS pathology may not be simply attributed to actin misregulation. This is the first comprehensive biochemical comparison of the actin assembly capacity for the known ALS-associated profilin variants.

## Methods

### Reagents and materials

All reagents were obtained from Fisher Scientific (Waltham, MA) unless otherwise stated.

### Protein plasmids, purification, and handling

We used a modified pET protein expression vector pMW172 containing the DNA sequences of wild-type profilin or the Y6D, C71G, M114T, E117G, G118V, R88E, and H120E profilin mutants (described previously) (Henty-Ridilla et al., 2017). Additional plasmids containing the DNA sequences flanked by NdeI and EcoRI sites for the A20T, T109M, R136W, and Q139L variants were synthesized and inserted into this same vector by Genscript (Piscataway, NJ). Profilin constructs (wild-type or ALS-variants) were transformed and expressed in Rosetta2(DE3) pRare2 (MilliporeSigma, Burlington, MA) competent cells. Cells were grown in Terrific Broth to OD_600_ = 0.6 at 37 °C, then induced with IPTG for 18 h at 18 °C. Cell pellets were collected by centrifugation and stored at -80 °C until purification. Profilin cell pellets were resuspended in 50 mM Tris HCl (pH 8.0), 1 mg/mL DNase I, 20 mg/mL PMSF, and 1× homemade protease inhibitor cocktail (0.5 mg/mL leupeptin, 1000 U/mL aprotinin, 0.5 mg/mL pepstatin A, 0.5 mg/mL antipain, 0.5 mg/mL chymostatin). Cells were incubated with 1 mg/mL lysozyme for 30 min and then lysed on ice with a probe sonicator at 100 mW for 90 s. The cell lysate was clarified by centrifugation at 278,000 × *g*. The supernatant was passed over a QHighTrap column (Cytiva, Marlborough, MA) equilibrated in 50 mM Tris-HCl (pH 8.0). Profilin was collected in the flow through and then applied to a Superdex 75 (10/300) gel filtration column (Cytiva) equilibrated in 50 mM Tris (pH 8.0), 50 mM KCl. Fractions containing profilin were pooled, aliquoted, snap-frozen in liquid nitrogen, and stored at -80 °C.

A modified pET23b vector containing the DNA sequence of either a constitutively active fragment of the formin mDia1 (mDia1 FH1-C) (amino acids 571-1255) or GFP-Tβ4 (Pimm et al., 2021) was transformed, induced, and collected as described for profilin (above). Cell pellets were resuspended in lysis buffer (2× PBS (pH 8.0) (2.8 M NaCl, 50 mM KCl, 200 mM sodium dibasic, 35 mM potassium monobasic), 20 mM imidazole (pH 7.4), 500 mM NaCl, 0.1% Triton-X 100, 14 mM BME) and lysed. Lysate was clarified via centrifugation for 30 min at 20,000 x *g* and the supernatant was flowed over cobalt affinity columns (Cytiva) equilibrated in low imidazole buffer (1× PBS (pH 8.0) supplemented with 20 mM imidazole (pH 7.4), 500 mM NaCl, 0.1% Triton-X 100, 14 mM BME). The 6×His-tagged proteins were eluted using a linear gradient into high imidazole buffer (1× PBS (pH 8.0) supplemented with 300 mM Imidazole (pH 7.4), 150 mM NaCl, 0.1% Triton-X 100, 14 mM BME) and the tag was cleaved with 5 mg/mL ULP1 protease for 2 h at 4 °C. proteins were concentrated in an ultrafiltration centrifugation device with an appropriate MWCO (4k MWCO for GFP-Tβ4 or 50k MWCO for mDia1(FH1-C)). GFP-Tβ4 concentrate was applied to a Superdex 75 (10/300) gel filtration column (Cytiva) equilibrated with gel filtration buffer (1× PBS (pH 8.0) supplemented with 150 mM NaCl, 14 mM BME). The mDia1(FH1-C) concentrate was applied to a Superose 6 Increase (10/300) gel filtration column (Cytiva) equilibrated with HEKG_5_ buffer (20 mM HEPES (pH 7.5), 1 mM EDTA, 50 mM KCl, 5% glycerol 14 mM BME). Fractions containing the pure proteins were pooled, aliquoted, snap-frozen in liquid nitrogen, and stored at -80 °C.

Rabbit skeletal muscle actin (RMA), Oregon-Green (OG) labeled-actin, N-(1-pyrenyl)iodoacetamide (pyrene) actin, and Alexa-488 actin were purified from acetone powder and labeled as described in detail (Cooper et al., 1984; Henty-Ridilla et al., 2017; Pimm et al., 2021; Spudich and Watt, 1971). Biotin-labeled actin was purchased from Cytoskeleton, Inc (Denver, CO). Fluorescently labeled actins and unlabeled actin were stored at -20 in 50% glycerol or -80 °C in G-buffer (3 mM Tris (pH 8.0), 0.5 mM DTT, 0.2 mM ATP, 0.1 mM CaCl_2_), respectively. Before use these proteins were dialyzed against freshly made G-buffer and pre-cleared at 279,000 × *g* to remove any polymer, nuclei, or aggregates.

All protein concentrations were determined by band densitometry from Coomassie-stained SDS-PAGE gels compared to a BSA standard curve. Gels were quantified using a LI-COR Odyssey imaging system (LI-COR Biotechnology, Lincoln, NE) and FIJI ImageJ software (Schindelin et al., 2012). Labeling stoichiometries were determined using the spectroscopy, molar extinction coefficients, and predetermined correction factors, as follows: unlabeled actin ε_290_ = 25,974 M^-1^cm^-1^, Oregon Green ε_496_ = 70,000 M^-1^cm^-1^, Pyrene actin ε_339_ = 26,000 M^-1^cm^-1^, Alexa-488 ε_495_ = 71,000 M^-1^cm^-1^. The correction factor used for Oregon Green was 0.12 and for Alexa-488 was 0.11. No correction factor was used for the pyrene label.

### Profilin binding assays

Profilin-actin binding experiments were performed by fluorescence polarization in binding mix (1× PBS (pH 8.0) supplemented with 150 mM NaCl). The binding constants of profilin (wild-type or ALS-variant) for actin monomers were determined in competitive binding assays with 10 nM unlabeled actin and 10 nM GFP-Tβ4 with variable concentrations of unlabeled profilin (to avoid potential interference with tags and binding partners and maintain the monomeric form of actin). Reactions were incubated at room temperature for 15 min and polarization was determined by exciting at 440 nm and measuring emission intensity at 510 nm with bandwidths set to 20 nm using a plate reader equipped with a monochromator (Tecan, Männedorf, Switzerland). Displacement of GFP-Tβ4 from actin monomers results in a decrease in polarization. Thus, the data were inverted (Y = -Y) to show profilin binding.

Profilin-poly-*L*-proline (PLP) binding constants were determined in direct binding assays with 10 nM unlabeled profilin (wild-type or mutant) and increasing concentrations of a FITC-mDia1-derived peptide (IPPPPPLPGVASIPPPP-PLPG), synthesized and labeled by Genscript and described previously (Kursula et al., 2008). Reactions were incubated at room temperature for 15 min and polarization was determined by exciting at 492 nm and measuring emission intensity at 520 nm with bandwidths set to 20 nm.

Resulting binding constants are the mean k_D_ produced from triplicate experiments. Non-linear curve fits for polarization experiments were performed using data normalized so that the smallest mean in each data set was defined as zero. Data were fit to the following curve using least squares regression with no constraints: Y = Y_0_-B_max_*(X/(K_D_+X)). Profilin proteins were pre-cleared at 279,000 × *g* before use in experiments.

### Bulk actin assembly assays

Bulk actin assembly assays were performed by combining freshly recycled 2 μM monomeric Mg-ATP actin (5% pyrene labeled), proteins or control buffers, and initiation mix (2 mM MgCl_2_, 0.5 mM ATP, 50 mM KCl). Total fluorescence was monitored using excitation 365 nm and emission 407 nm in a Tecan plate reader. Reactions for each replicate were performed on the same plate. Reactions were initiated by adding actin to reactions using a multichannel pipette. Mean values shown in Figure S1 were averaged from three independent experiments.

### Total internal reflection fluorescence (TIRF) microscopy assays

TIRF microscopy chambers with PEG-silane coated coverslips were prepared as previously described (Henty-Ridilla et al., 2017, 2016; Pimm et al., 2021; Smith et al., 2013). Chambers were conditioned with the following reagents and order: 1% BSA, 4 μg/mL streptavidin, 1% BSA, and then 1× TIRF buffer supplemented with 0.25% methylcellulose [4000 cP] (20 mM imidazole (pH 7.4) 50 mM KCl, 1 mM MgCl_2_, 1 mM EGTA, 0.2 mM ATP, 10 mM DTT, 40 mM glucose). A DMi8 inverted microscope equipped with 120-150 mW solid-state lasers, a 100× Plan Apo 1.47 N.A. oil-immersion TIRF objective (Leica Microsystems, Wetzlar, Germany), and an iXon Life 897 EMCCD camera (Andor; Belfast, Northern Ireland) was used for all TIRF experiments. Time-lapse movies were acquired at 5 s intervals with 50 ms exposure, 488 nm excitation, 4% laser power for experiments with Oregon-green or Alexa-488 actin. Reactions were introduced into the flow chamber using a Luer slip system and a syringe pump (Harvard Apparatus, Holliston, MA). All reactions are timed from the initial mixing of proteins rather than the start of image acquisition (usually delayed by 15-20 s). Actin was loosely tethered to the cover glass surface using an avidin-biotin conjugation system. Actin nucleation was measured as the number of actin filaments present 100 s or 240 s after the initiation of the reaction. Actin filament elongation rates were measured as the slope of a line generated from the length (μm) of actin filaments over time for at least four consecutive movie frames. Rates were multiplied by a correction factor of 370 (the number of actin subunits per micron filament) (Pollard et al., 2000). Background subtraction of raw TIRF images was achieved using a 50 pixel rolling-ball radius in FIJI software.

For urea chemical denaturation experiments we exposed stock concentrations of profilin (wild-type or ALS-variant) to 5 M urea for 30 min to denature the proteins. To facilitate refolding, the concentration of urea was reduced to 344 mM in TIRF buffer. Thus, the final concentration of urea was 316.5 μM once actin was added to initiate the reaction. This final urea concentration was used to keep a consistent 5 μM concentration of all profilins. After a 30 min refolding period, urea-treated profilins were assessed for actin polymerization activities in TIRF assays (as described above). The final concentration of urea in all conditions (including actin controls) was 344 mM.

### Data analysis, statistics, and presentation

GraphPad Prism (version 9.3.1) (GraphPad Software, San Diego, CA) was used for all data analyses and to perform all statistical tests. The specific details for each experimental design, sample size, and specific statistical tests are available in the figure legends. Power analysis was performed for all datasets and all datasets were tested for normality before performing t-tests or ANOVA. P-values lower than 0.05 were considered significant for all analyses. PyMOL (version 2.3.4) (Schrödinger, Inc, New York, NY) was used for molecular visualizations. Adobe Illustrator (version 2021 25.4.1) (Adobe, San Jose, CA) was used for data presentation. Datasets for each figure have been deposited in Zenodo and are available with request at: https://doi.org/10.5281/zenodo.5975762.

## Results & Conclusions

### ALS-linked profilin proteins bind actin monomers with different affinities

There are eight ALS-associated mutations in profilin (A20T, C71G, T109M, M114T, E117G, G118V, R136W, and Q139L) (Castellanos-Montiel et al., 2020; Gregory et al., 2020). The C71G, E117G, G118V mutations are located within or directly adjacent to the actin binding surface of profilin (Figure 1A) (Ferron et al., 2007). The A20T, T109M, R136W, and Q139L mutations reside on or adjacent to the surface of profilin responsible for interactions with actin nucleation ligands that contain poly-*L*-proline (PLP) sequences i.e., formins or Ena/VASP (Figure 1A) (Ferron et al., 2007). The M114, E117, and G118 residues are also associated with microtubule regulation and mediate competitive binding interactions for profilin between actin monomers and microtubules (Figure 1A) (Henty-Ridilla et al., 2017). The close proximity to the actin binding surface on profilin and the essential and well characterized roles for profilin in regulating actin dynamics have led several groups to hypothesize that ALS-linked variants of profilin perturb actin regulation. However, the extent of direct actin binding for each ALS-associated mutation in profilin has not been determined.

**Fig. 1.**
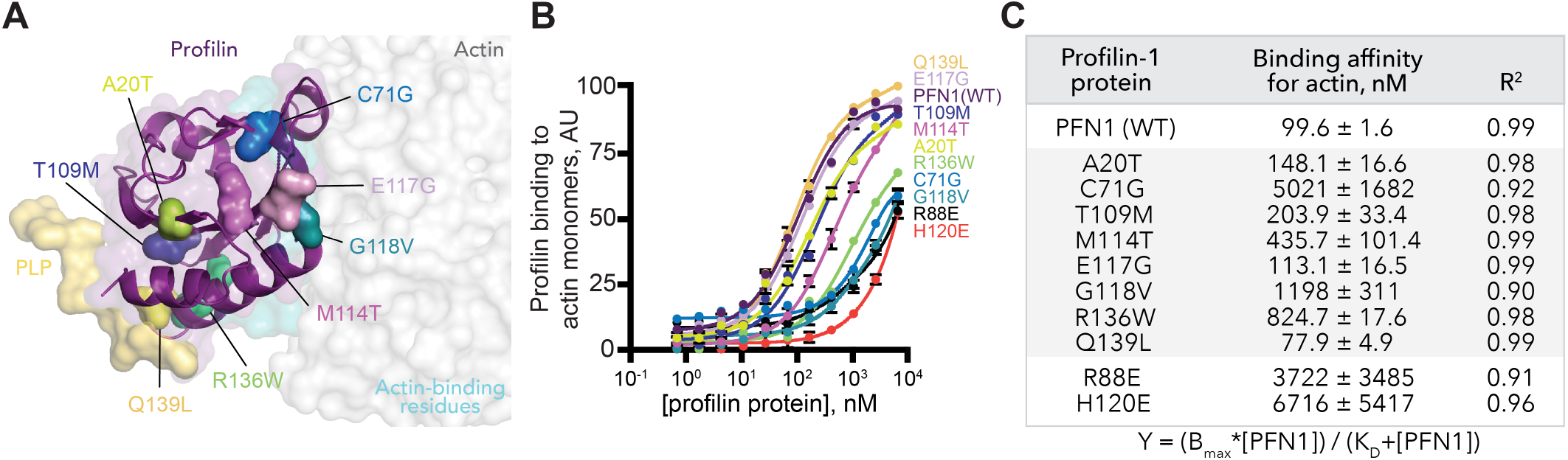
ALS-associated profilin variants vary in actin monomer binding capacity. **(A)** View of profilin (purple) simultaneously bound to an actin monomer (gray) and a poly-*L*-proline (PLP) peptide from VASP (yellow). View was modeled using PDB: 2BTF (Schutt et al., 1993) and PDB: 2PAV (Ferron et al., 2007). Profilin surfaces contacting actin are highlighted (cyan). The positions of eight ALS-relevant residues on profiin-1: A20T (chartreuse), C71G (cerulean), T109M (dark blue), M114T (magenta), E117G (lavender), G118V (teal), R136W (green), and Q139L (gold) are indicated. **(B)** Competitive fluorescence polarization measurements of 10 nM GFP-thymosin β4 (GFP-Tβ4), 10 nM unlabeled actin monomers, and increasing concentrations of profilin (wild-type (PFN1) or ALS-variant). Curves shown are the mean of three separate experiments. Error bars indicate SEM. **(C)** Table indicating the binding affinities for profilin (wild-type or ALS-variant) determined from (B).

We recombinantly expressed and purified untagged versions of profilin, the eight ALS-associated profilin proteins, and two previously characterized profilin controls deficient for actin binding (R88E and H120E) (Ezezika et al., 2009; Kovar et al., 2006; Lu and Pollard, 2001; Suetsugu et al., 1998). To determine and compare the binding affinities for wild-type profilin and each ALS-linked profilin for actin monomers, we monitored fluorescence polarization in competitive binding assays with GFP-thymosin β4 (Figures 1B and 1C). Wild-type profilin bound actin monomers with k_D_ = 99.6 nM ± 1.6, consistent with previous studies (Figures 1B and 1C) (Aguda et al., 2006; Goldschmidt-Clermont et al., 1992; Pimm et al., 2021). Profilin proteins harboring the A20T (k_D_ = 148.1 nM ± 16.6), T109M (k_D_ = 203.9 nM ± 33.4), or E117G (k_D_ = 113.1 nM ± 16.5) mutations each bound actin monomers with similar affinities as wild-type profilin (Figure 1B and 1C). One variant (Q139L) bound actin monomers 1.3-fold more efficiently than wild-type (k_D_ = 77.9 nM ± 4.9). However, the majority of ALS-associated profilins displayed significantly weaker affinities for actin monomers (Figures 1B and 1C). G118V (k_D_ = 1,198 nM ± 311) and R136W (k_D_ = 824.7 nM ± 17.6) bound actin monomers ∼ 10-fold less efficiently. The C71G variant is the most impaired for actin monomer binding (k_D_ = 5,021 nM ± 1682), with an affinity similar to the R88E (k_D_ = 3,722 nM ± 3,485) and H120E (k_D_ = 6,716 nM ± 5,417) actin-binding impaired controls (Figure 1C).

Our results demonstrate that each of the ALS-associated mutations in profilin influence the affinity of profilin for actin monomers. This result is anticipated for the C71G, E117G, G118V mutations due to their close proximity to the surface the actin binding surface on profilin (Figure 1A). The differences in actin monomer affinities for M114T, R136W, and Q139L are less clear as these residues are not adjacent to the actin-binding surface and some are located on the opposite side of the molecule (Figure 1A). Previous characterizations of profilin harboring the M114T, G118V, or T109M mutations imply that these variants have subtle folding stability defects (Boopathy et al., 2015; Freischmidt et al., 2015). Similar properties may distort the profilin-actin interaction surface for other profilin variants to produce these distinct affinities.

### ALS-linked profilin variants reduce actin assembly in vitro

Related to its ability to bind to actin monomers, profilin sterically suppresses the formation of new actin filaments (Blanchoin et al., 2014; Skruber et al., 2020). To assess this concept further, we compared the average rates of actin assembly in the presence of each ALS-associated profilin variant in bulk pyrene actin-fluorescence assays (Figure S1A). Compared to the actin alone control, less total actin polymer was made in the presence of profilin (Figure S1A). Actin filaments formed in the presence of A20T, G118V, E117G, or T109M polymerized to similar levels as wild-type profilin (Figure S1). Reactions containing actin and C71G, M114T, or R136W had levels of actin polymerization above wild-type profilin but were still able to suppress bulk filament assembly compared to reactions lacking profilin (Figure S1A). Q139L most strongly inhibited actin filament polymerization in these assays (Figure S1A). These results suggest that profilin binding affinities generally correlate with the actin nucleation strength of each ALS-related profilin.

Actin assembly consists of two parameters, filament nucleation (total number of filaments) and the rate of filament elongation (incorporation of monomers into polymerizing filaments). Bulk actin pyrene assays measure both of these parameters but are heavily influenced by filament nucleation, rather than elongation (Rosenbloom et al., 2021; Zweifel et al., 2021). To distinguish these polymerization phases more carefully, we used total internal reflection fluorescence (TIRF) microscopy to monitor actin assembly in the presence of each ALS-linked profilin (Figure 2). Consistent with profilin-actin monomer binding affinities and bulk fluorescence assays, TIRF reactions containing 5 μM profilin (wild-type or ALS-mutants) contained fewer filaments than reactions containing 1 μM actin alone (Figures 2A and 2B). Specifically, reactions performed in the presence of A20T, T109M, or E117G had mean filament numbers similar to wild-type profilin, whereas profilins with weaker binding affinities (M114T, G118V, R136W) had elevated means (Figures 2A and 2B). Congruent with prior analyses, C71G had the highest mean actin filament nucleation counts, reaching similar levels as the R88E and H120E mutations in profilin that result in reduced monomer binding (Figures 2A, 2B, S2A, and S2B). In addition, reactions containing Q139L very strongly inhibited filament nucleation (Figures 2A and 2B). Importantly, these nucleation trends occur over many concentrations of profilin (wild-type or ALS-related mutants), with the exception of proteins profoundly impaired for actin binding (i.e., C71G or the R88E and H120E controls) (Figures S3A-C). This demonstrates that most of these profilin variants retain actin functions regardless of any variation that may arise from protein folding stability.

**Fig. 2.**
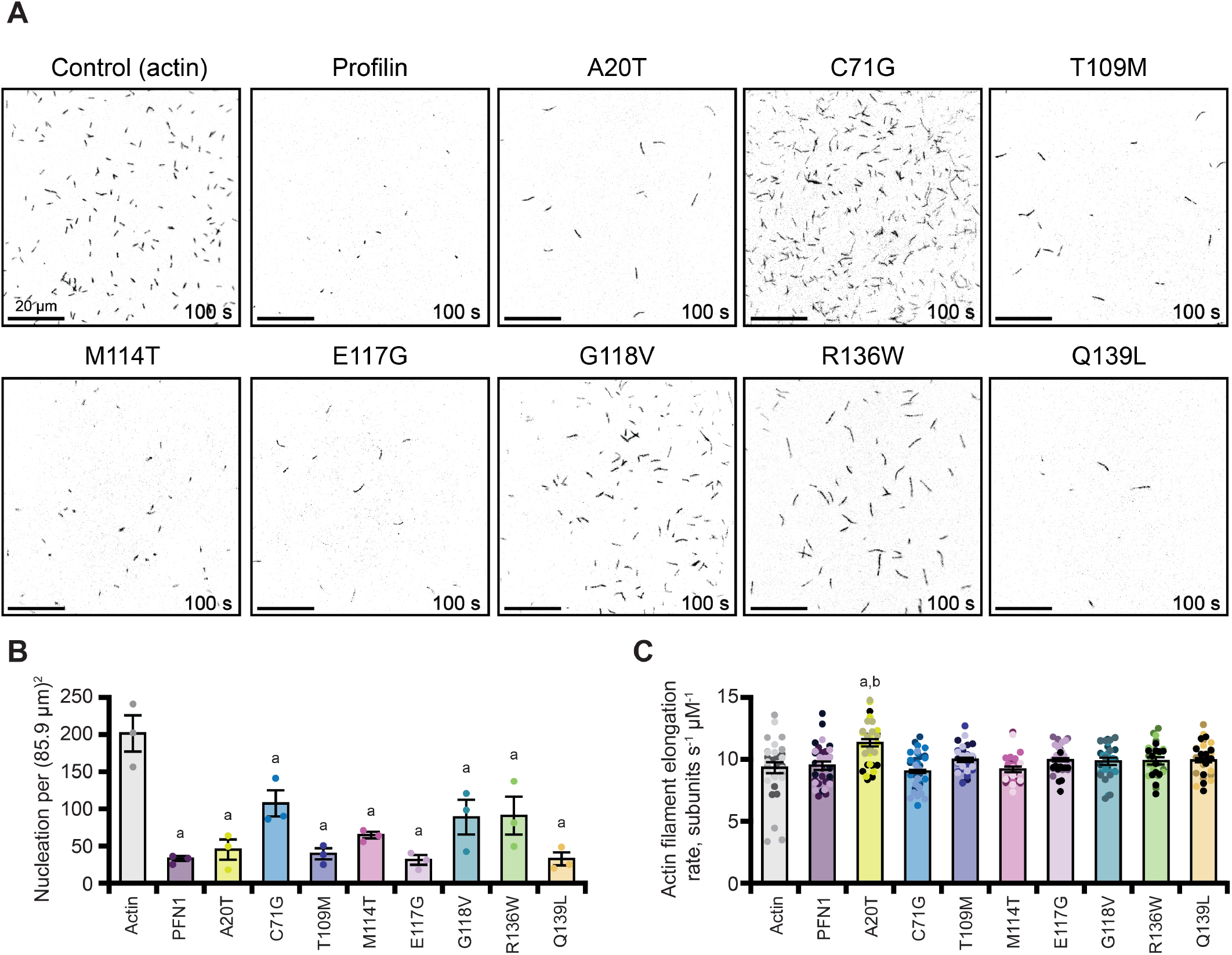
Profilin variants inhibit actin filament nucleation. **(A)** Representative fields of view (FOV) from TIRF microscopy assays containing 1 μM actin (10% Oregon Green (OG)-labeled, 0.6 nM biotin-actin) assembled in the absence (control) or presence of 5 μM profilin (wild-type or mutant). Scale bars, 20 μm. **(B)** Mean count of actin filaments visible from TIRF reactions shown in (A). Measurements were made 100 s after the actin polymerization was initiated. Each dot represents counts from different experimental replicates. **(C)** Distribution of actin filament elongation rates from TIRF reactions as in (A). Dots represent individual measurements (n = 11 per replicate or n = 33 total). Shaded values show the distribution of different independent experimental replicates (n = 3). Error bars indicate SEM. a, significantly different (P < 0.05) than actin control; b, significantly different (P < 0.05) than wild-type profilin control. Number of measurements determined by power analysis. Significant differences for nucleation experiments were determined by one-way ANOVA with Bonferroni post-hoc analysis. For elongation rate experiments, Tukey post-hoc analysis was used.

Profilin has spectacular differences in affinities for actin monomers (k_D_ = ∼ 100 nM) compared to filament ends (k_D_ = 225 μM) (Courtemanche and Pollard, 2013; Funk et al., 2019; Pernier et al., 2016; Zweifel et al., 2021; Zweifel and Courtemanche, 2020). Thus, profilin avoids interference with active actin polymerization by rapidly disassociating from the ends of growing actin filaments (Courtemanche and Pollard, 2013; Funk et al., 2019; Jégou et al., 2011). To assess whether the ALS-linked profilins contribute to the filament elongation phase of actin assembly we measured elongation rates of actin filaments from TIRF movies. In the presence of profilin, the mean rate of actin filament polymerization was 9.5 ± 0.3 (SEM) subunits s^-1^ μM^-1^, which is not significantly different from the rate of 9.3 ± 0.4 subunits s^-1^ μM^-1^ measured for actin filaments in reactions without profilin or the R88E and H120E negative controls (Figures 2C and S2C). Apart from A20T, actin filaments polymerized in the presence of each ALS-linked profilin displayed similar behaviors as wild-type profilin with elongation rates ranging from 9.1–10.1 subunits s^-1^ μM^-1^ (Figure 2C). The polymerization rate of actin filaments in the presence of A20T was significantly elevated (11.4 ± 0.3 subunits s^-1^ μM^-1^) compared to wild-type profilin (Figure 2C). This effect is surprising because the A20T variant has lower affinity for actin monomers in both binding and nucleation experiments, and is located opposite to the actin-binding surface of profilin. This may indicate that this mutation promotes faster disassociation of profilin from actin (monomers or filament ends) or that the conformation of this profilin deviates from wild-type. Thus, the ALS-related variants are most disruptive to the nucleation phase of actin assembly.

### Chemical denaturation further destabilizes ALS-linked profilin variants and alters actin polymerization activities

The C71G, T109M, M114T and G118V mutations subtly destabilize the tertiary structure of profilin (Boopathy et al., 2015; Freischmidt et al., 2015; Schmidt et al., 2021). These effects are thought to impede normal profilin functions in actin assembly but have not been assessed or compared across all identified ALS-associated variants. Wild-type profilin has a very robust structure with efficient folding dynamics and can refold after denaturization with high molar urea (Kaiser et al., 1989; Krishnan and Moens, 2009). To reiterate, we purified profilin (wild-type and variants) without chemical denaturation and using a tag-less purification scheme. To compare the actin polymerizing activities of profilin (wild-type or ALS-variant) following chemical destabilization and refolding, we exposed profilin (wild-type or ALS-variant) to 5 M urea for 30 min to denature the protein. To facilitate protein refolding, the concentration of urea was reduced to 344 mM in buffer (50 mM Tris (pH 8.0), 50 mM KCl). After a 30 min refolding period, urea-treated profilins were assessed for actin polymerization activities in TIRF assays (Figure 3). Compared to untreated controls (Figure 2), fewer actin filaments were nucleated in the presence of this concentration of urea and no negative effects were observed for actin filament elongation (9.2 ± 0.5; Figure 3). Similar to profilin positive controls, we observed fewer actin filaments nucleated and no effects to actin filament elongation for actin-binding impaired negative controls (R88E and H120E) (Figures S4A-C).

**Fig. 3.**
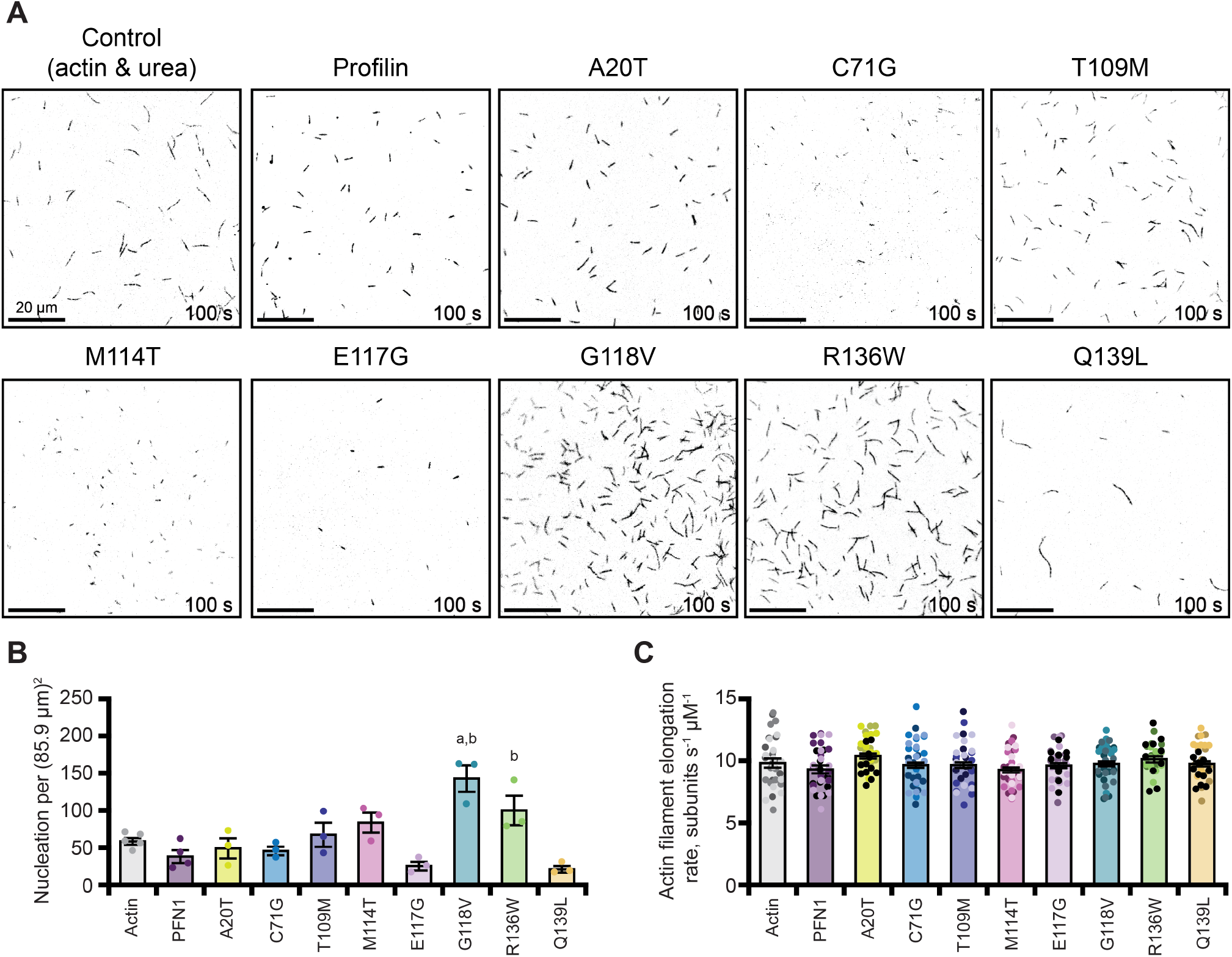
Urea denaturation influences profilin variant stability and further alters actin polymerization activities. **(A)** Representative fields of view (FOV) from TIRF microscopy assays containing 1 μM actin (10% Alexa-488 labeled, 0.6 nM biotin-actin) in the absence (control) or presence of 5 μM profilin (wild-type or variant) that was treated with 5 M urea for 30 min, diluted to 344 mM in TIRF buffer, and allowed to refold for 30 min. After the refolding period urea-treated profilins were assessed for actin activities in TIRF assays. The final concentration of urea in each of these experiments (including actin controls) was 316.5 mM. Scale bars, 20 μm. **(B)** Mean count of actin filaments as in (A) 100 s after actin polymerization was initiated. Each dot represents counts from independent experimental replicates. **(C)** Distribution of actin filament elongation rates from TIRF reactions as in (A). Dots represent individual measurements (n = 11 per replicate or n = 33 total). Shaded values show the distribution of different independent experimental replicates (n = 3). Error bars indicate SEM. a, significantly different (P < 0.05) than actin control; b, significantly different (P < 0.05) than wild-type profilin control. Number of measurements determined by power analysis. Significant differences for nucleation experiments were determined by one-way ANOVA with Bonferroni post-hoc analysis. For elongation rate experiments, Tukey post-hoc analysis was used.

In the presence of urea-treated profilin, actin filament nucleation was suppressed albeit to lower levels than reactions performed with untreated protein (Figures 3A, 3B, 2A, and 2B). Similar to wild-type profilin, the nucleation trends for several of the ALS-related variants are the same as the untreated proteins: each variant suppresses actin filament nucleation to some extent (Figures 3A and 3B). However, T109M, M114T, G118V, and R136W display elevated levels of actin nucleation compared to untreated controls. Consistent with other reports for some of these proteins, this may indicate that these variants do not fold as efficiently as the wild-type protein (Boopathy et al., 2015; Figley et al., 2014). Urea treatment of the already impaired C71G variant tends to have fewer actin filaments than controls (Figures 3A and 3B). This may reinforce evidence that this variant is unstable or that it forms polymerization incompetent complexes with actin. Urea treatments did not influence the elongation phase of actin polymerization for any of the variants tested, except for A20T which no longer displayed an elevated mean elongation rate compared to wild-type profilin (Figure 3C). Regardless of the residue position (i.e., closer or further from the actin binding surface), it is difficult to draw strong conclusions about profilin folding stability and actin polymerization. Notably, refolding trends do not seem to affect all profilin variants equally. The range of observed stability effects (here and elsewhere) may be influenced by purification schemes or protein handling.

### Formin-mediated actin polymerization is influenced by ALS-linked profilin variants

ALS-associated profilin variants clearly influence actin assembly through direct binding interactions with actin monomers. However, the indirect role of profilin with actin polymerization stimulating ligands like formin or Ena/VASP is arguably the most influential contribution for profilin in a cellular context (Harker et al., 2019; Pimm et al., 2020; Pimm and Henty-Ridilla, 2021; Rotty et al., 2015; Skruber et al., 2020; Suarez et al., 2015). Profilin can simultaneously bind actin and these or other poly-L-proline (PLP)-containing ligands (Ferron et al., 2007). For formin proteins, interactions between profilin-bound actin and PLP sites in the formin homology 1 (FH1) domains orient actin monomers to stimulate actin assembly (Courtemanche, 2018; Courtemanche and Pollard, 2012; Zweifel and Courtemanche, 2020). Thus, we sought to assess the effects of each ALS-associated profilin variant on formin-mediated actin polymerization. For these analyses we chose the representative formin mDia1 for three reasons: 1) mDia1 polymerizes actin at the highest rates recorded among all the mammalian formins, thus impairments might be easier to detect; 2) wild-type profilin and mDia1 interact at discrete sites of cellular actin polymerization (Jacquemet et al., 2019); and 3) mDia1 is known to interact with a subset of ALS-associated profilins (G118V and M114T) (Schmidt et al., 2021).

First, we compared the average rates of formin-mediated actin assembly in the presence of each ALS-associated profilin in bulk pyrene fluorescence assays (Figure S1B). As expected, compared to the control (actin and mDia1), less average actin assembly occurs in the presence of wild-type profilin (Figure S1B). As expected, the total actin polymerization measured is less, as these reactions contain longer dimmer actin filaments due to the labeling procedure of actin at cysteine 374 (Rosenbloom et al., 2021; Zweifel et al., 2021). The ALS-associated profilin variants group into three different classes: reactions that appear similar to actin alone, reactions that polymerize to similar levels as wild-type profilin, and reactions with very little polymerization (Figure S1B). Reactions performed in the presence of A20T, C71G, or M114T appear similar to reactions lacking profilin (Figure S1B, right). In agreement with these findings, these mutants exhibit reduced ability to bind actin monomers and suppress actin filament nucleation in reactions lacking formin (Figures 1B, 1C, and 2B). Reactions containing E117G, G118V, or T109M appear to stimulate formin-based actin assembly but have less total polymerization compared to control reactions containing wild-type profilin (Figure S1B, left). Lastly, reactions containing R136W or Q139L have markedly less actin polymerization compared to wild-type profilin (Figure S1B, left). Neither of these profilin variants appear to stimulate formin-based actin assembly at all. This result is interesting as the R136W variant may be separating formin-profilin and profilin-actin functions, whereas the Q139L variant appears to suppress actin filament nucleation and elongation with actin alone or formin. In sum, rather than direct defects in actin monomer binding, the most striking deficiencies to profilin-actin assembly manifest in formin-mediated actin assembly.

To investigate these observations more carefully, we performed TIRF microscopy to evaluate the detailed assembly of single actin filaments in the presence of each ALS-related profilin variant and constitutively active mDia1 (Figure 4A). As expected and consistent with past reports, the reactions containing 10 nM mDia1(FH1-C) and actin display enhanced actin filament nucleation compared to reactions lacking formin (Figures 2A, 2B, 4A, and 4B) (Breitsprecher et al., 2012; Breitsprecher and Goode, 2013; Chesarone et al., 2010; Henty-Ridilla et al., 2016). The actin-binding impaired negative controls (R88E or H120E) did not suppress formin-mediated actin nucleation (Figure S5A and S5B). The general distribution of actin filament nucleation efficiencies is similar to reactions without mDia1(FH1-C), just at higher, formin-stimulated, levels (Figures 2B and 4B). C71G, M114T, and R136W deviate from this trend (Figures 4B and 2B). Reactions performed with C71G or R136W display a reduced mean in filament nucleation compared to reactions lacking formin (Figures 4B and 2B). Reactions containing actin, formin, and M114T nucleate actin filaments at rates significantly different from wild-type profilin or control conditions that lack profilin (Figure 4B). This may indicate that combinatory effects between M114T and formin affect the nucleation phase of actin assembly. In sum, the general trends of formin-based actin filament nucleation are similar to reactions containing wild-type profilin for all ALS-associated variants (albeit at higher levels) with the exception of R136W and C71G. Somehow R136W is suppressing actin polymerization in the presence of formin more than its absence. One possible explanation for this result could be that this variant binds to specific PLP tracks more tightly, stalling efficient elongation or effectively blocking formin-based assembly.

**Fig. 4.**
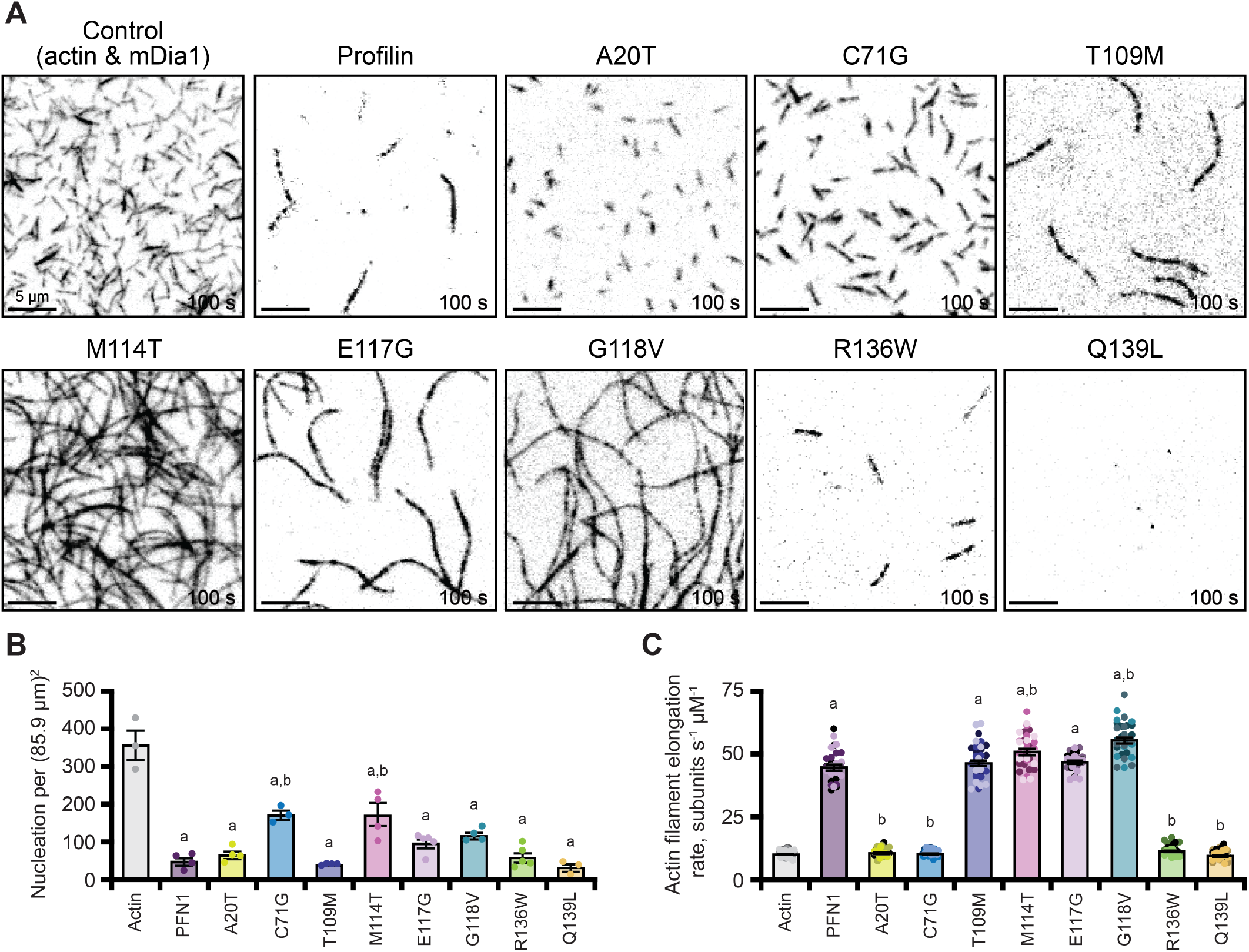
Several profilin variants fail to stimulate formin-mediated actin polymerization. **(A)** Representative time lapse TIRF images from actin polymerization assays in the presence of formin and absence or presence of profilin. Reactions contain 1 μM actin (10% Alexa-488 labeled, 0.6 nM biotin-actin) and 10 nM mDia1(FH1-C) without or with 5 μM profilin (wild-type or ALS mutant as labeled). **(B)** Graphical representation of actin filament nucleation for TIRF movies as in (A); individual data points represent the number of filaments per field of view at 100 s after initiation of actin assembly from separate polymerization experiments. Error bars, represent SEM. **(C)** Distribution of actin filament elongation rates from TIRF reactions as in (A). Dots represent individual measurements (n = 11 per replicate or n = 33 total). Shaded values show the distribution of different independent experimental replicates (n = 3). Error bars indicate SEM. a, significantly different (P < 0.05) than actin control; b, significantly different (P < 0.05) than wild-type profilin control. Number of measurements determined by power analysis. Significant differences for nucleation experiments were determined by one-way ANOVA with Bonferroni post-hoc analysis. For elongation rate experiments, Tukey post-hoc analysis was used.

We next assessed whether the ALS-linked variants in profilin influenced the elongation phase of formin-mediated actin assembly in TIRF assays (Figure 4C). Consistent with previous reports, the addition of mDia1 to reactions containing wild-type profilin stimulated actin assembly from 10.2 ± 0.23 sub-units s^-1^ μM^-1^ (actin and formin alone) to 44.6 ± 1.0 subunits s^-1^ μM^-1^ (actin, formin, and profilin) (Figure 4C) (Henty-Ridilla et al., 2016; Kovar et al., 2006; Pimm et al., 2021). As expected, the actin-binding impaired negative control profilin mutants R88E and H120E did not stimulate actin filament elongation (Figures S5A and S5C). T109M and E117G stimulated formin-based actin assembly and were not significantly different than wild-type profilin (Figure 4C). Half of the profilin variants (A20T, C71G, R136W or Q139L) failed to stimulate formin-mediated actin assembly (Figure 4C). This result is somewhat expected for the A20T, R136W or Q139L variants, which lie in the PLP-interacting region of profilin that is required for formin interaction. The C71 residue lies on the opposite side of the profilin molecule. However, its lack in ability to stimulate formin-based assembly may be explained by its general folding instability (Figure 3) (Boopathy et al., 2015; Schmidt et al., 2021). This observation may conflict with a recent report showing this variant was still capable of stimulating formin polymerization at reduced levels in different TIRF conditions that produce tension/pulling forces on growing actin filaments (Schmidt et al., 2021).

Intriguingly, two ALS-associated profilin variants, G118V and M114T, displayed enhanced formin-stimulated actin assembly (Figure 3B). Each of these variants have weakened affinities for actin monomers (Figures 1B and 1C). Bulk pyrene assays did not reveal these elongation-based phenotypes, indicating that these profilins may have an even stronger preference than wild-type profilin for unlabeled actin monomers. Explanations for the observed stimulation of formin-mediated actin assembly include the possibility that these mutants have a higher off-rate for actin monomers, or a change in the affinity for any of the fifteen specific PLP motifs present in the FH1 domain of mDia1 (Courtemanche, 2018; Zweifel and Courtemanche, 2020). In summary, while actin binding is important for profilin function, the most striking defects associated with the ALS-associated profilin variants appear with formin. Further, the ALS-associated profilin variants differentially disrupt diverse facets of profilin function in actin assembly (i.e., monomer binding, nucleation, and/or formin-based elongation).

### ALS-associated profilin variants bind mDia1 PLP with similar affinities

Half of the ALS-related profilin variants failed to stimulate formin-mediated actin assembly. A simple mechanism to explain the loss of formin-mediated actin assembly for the A20T, C71G, R136W, and Q139L variants is weakened or lost affinity for binding to formin. To test this hypothesis, we performed fluorescence polarization with each profilin variant (Figure 5). For these experiments we used titrations of a previously characterized, >90% pure, FITC-conjugated peptide sequence containing two poly-*L*-proline (PLP) motifs from mDia1 (IPPPPPLPGVASIPPPPPLPG) (Kursula et al., 2008). This peptide sequence binds multiple profilin isoforms and contains the minimally required length for efficient profilin binding (Kursula et al., 2008). Wild-type profilin bound the mDia1 PLP peptide in a manner consistent with studies using similar peptides and constitutively active fragments containing the FH1 domains of mDia1 (k_D_ = 2.71 μM ± 0.11) (Figure 5) (Boopathy et al., 2015; Kursula et al., 2008; Schmidt et al., 2021). Both actin-binding deficient mutants R88E and H120E bound the mDia1 peptide with similar affinity as wild-type, however the PLP-deficient mutant Y6D did not bind the peptide at all (Figures 5A and 5B)(Ezezika et al 2009). Each ALS-associated profilin variant bound the peptide with relatively similar affinities ranging from 1.65 to 3.47 μM.

**Fig. 5.**
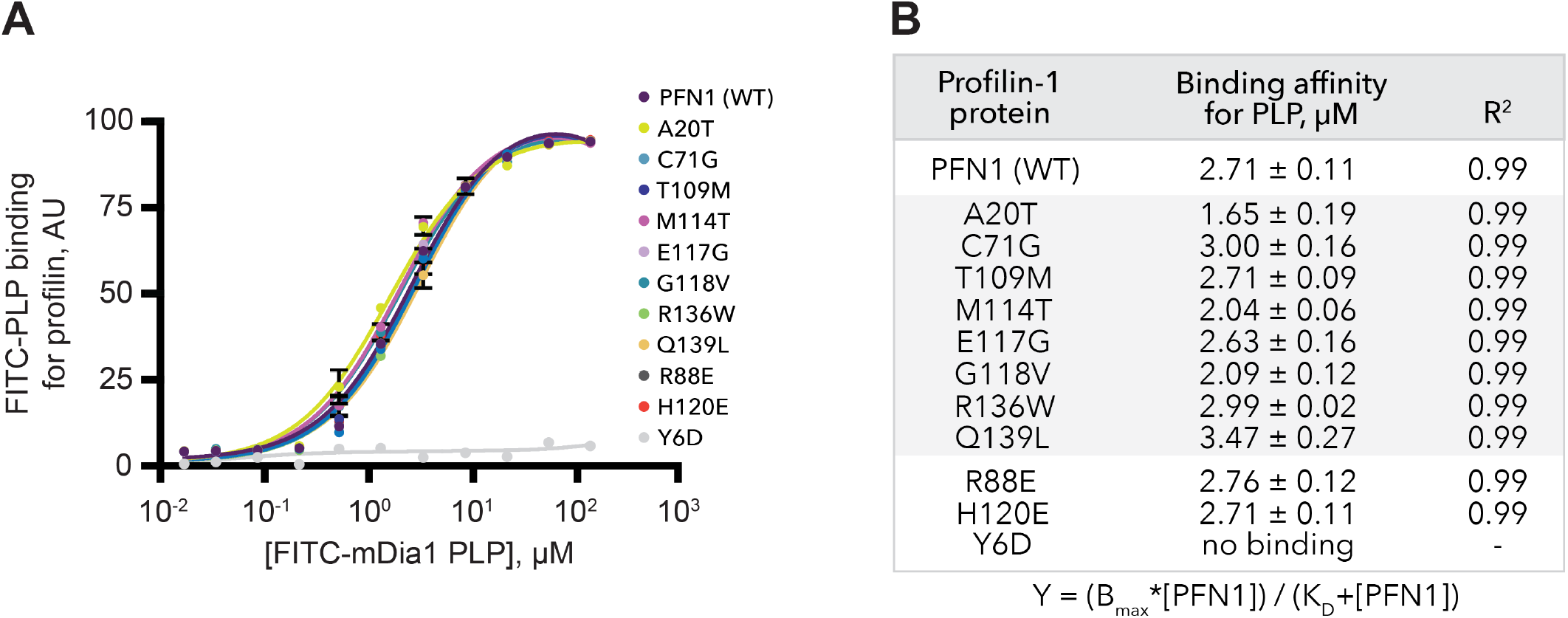
Profilin variants display similar binding affinities for an mDia1-based poly-*L*-proline peptide. **(A)** Fluorescence polarization measurements of 10 nM unlabeled profilin (wild-type or ALS mutant, as indicated) mixed with increasing concentrations of FITC-labeled mDia1-PLP peptide (IPPPPPLPGVASIPPPPPLPG, based on two consecutive profilin-binding domains of mDia1). Curves shown are the mean of three separate experiments. Error bars indicate SEM. Y6D was included as a PLP-binding deficient negative control (gray). **(B)** Profilin affinities for mDia1-PLP peptide from polarization data in (A); affinities (mean ± SEM calculated from the displayed equation) and R^2^ values are from three independent experiments. Y6D negative control (with additional wild-type profilin reference controls) was performed on a different plate and day than other proteins. There was no statistical difference between the wild-type profilin affinities for PLP between these plates so we included the Y6D control on the graph. Raw fluorescence intensities of the FITC peptide did not move out of the linear range for these analyses.

The Q139L variant bound this PLP peptide sequence the weakest. (k_D_ = 3.47 μM ± 0.27). Additional studies exploring other amino acid substitutions at this position suggest this residue is prone to instability (Del Poggetto et al., 2016; Muller et al., 2005). In contrast, the A20T variant, which also resides on the PLP-binding surface, bound the PLP peptide with higher affinity than wild-type (k_D_ = 1.65 μM ± 0.19) (Figures 1A, 5A, and 5B). How does a variant that binds PLP better fail to stimulate formin-based actin elongation? Perhaps this version of profilin blocks critical PLP binding sites required for the efficient transfer of actin monomers to growing actin filament ends or significantly decreases the off-rate of profilin-actin complexes from formin FH1 domains. One possible explanation is that the Q139L and A20T substitutions affect hydrogen bonding interactions at the profilin-PLP interface. In this scenario, Q139L subtracts one putative hydrogen bonding site and weakens the interaction, whereas the A20T substitution adds a possible hydrogen bonding site. Q139L also has a slightly higher affinity for actin, which may increase the overall dwell time of profilin bound to actin at the growing end of an actin filament.

Similar to other work, the M114T and G118V variants bound the mDia1-derived peptide with higher affinity than wild-type, k_D_ = 2.04 μM ± 0.06 and k_D_ = 2.09 μM ± 0.12, respectively. Each of these variants enhanced formin-mediated actin elongation. While both mutations reside far from the PLP binding surface (Figure 1A), they may enhance formin-based elongation by increasing the off-rate of profilin from PLP sequences in the formin FH1 domains (Figure 5A and 5B) (Schmidt et al., 2021). Alternatively, binding differences seen with this two PLP motif-containing peptide may be accentuated in formins that contain many PLP motifs; for example, mDia1 contains at least fifteen PLP binding sites on each of the two FH1s present in the active dimer (Courtemanche, 2018; Courtemanche and Pollard, 2012; Paul et al., 2008; Zweifel and Courtemanche, 2020). In conclusion, each of the ALS-associated profilin variants binds an mDia1-derived PLP peptide with similar affinities, despite varied capacities for stimulating formin-based actin assembly. Notably, the Y6D negative control does not bind the PLP peptide, consistent with previous reports (Ezezika et al 2009). However, these experiments do not assess whether the ALS-related profilin variants have more significant differences when binding other PLP-containing ligands including other formin proteins, Ena/VASP, survival motor neuron protein (SMN), or exportin-6.

## Discussion

The sum of ALS-associated defects to the actin cytoskeleton are often credited to mutations in the canonical actin monomer binding protein profilin. Profilin is an essential regulator of the neuronal cytoskeleton through many direct (i.e. binding actin monomers, tubulin dimers, or microtubules) and indirect mechanisms linked to cellular cofactors like formin, Ena/VASP, SMN, and exportin-6 (Figure 6A)(Bowerman et al., 2009; Henty-Ridilla et al., 2017; Murk et al., 2021; Pimm et al., 2021; Skruber et al., 2020; Stüven et al., 2003; Suarez and Kovar, 2016). However, the mechanisms defining the function of the ALS-associated mutations in actin assembly are challenging to interpret from cell-based studies and not well defined or quantitatively compared across all variants. To gain insight into the underlying mechanisms that define profilin-mediated ALS, we performed an in vitro study exploring the function of each of the eight ALS-related variants in actin assembly (summarized in Figure 6B). We learned that ALS-linked profilin proteins bind actin monomers with a broad range of affinities that generally correlate with the strength of actin filament nucleation for each variant (i.e., stronger binders inhibit nucleation) (Figures 1C and 2B). ALS-variants display a range of stability and folding defects. Here we show that further perturbation of protein folding can influence the ability of profilin to nucleate actin filaments (Figure 3B). We also observed the effects of each profilin variant on formin-mediated actin polymerization. The ALS-variants bind an mDia1-poly-*L*-proline (PLP) sequence with similar affinities yet vary in their ability to regulate formin actin assembly (Figures 4 and 5B) (Figure 6B).

**Fig. 6.**
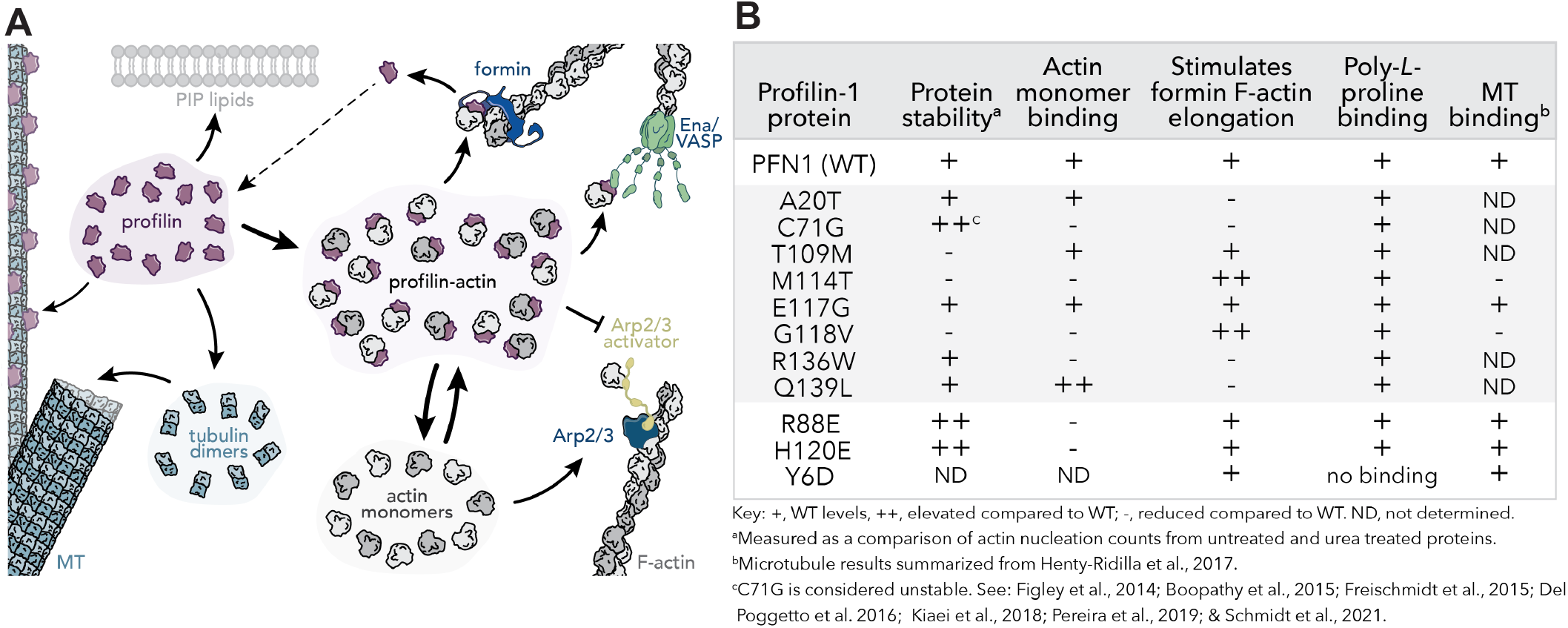
Summary of profilin protein activities. **(A)** Cartoon detailing the distribution of profilin for various ligands. PIP lipids, microtubules (MT), tubulin dimers, actin monomers, and several actin nucleation promoting factors (i.e., formin, Ena/VASP, the Arp2/3 complex) are each influenced by profilin-binding and disruptions elicited by the eight ALS-related profilin variants may contribute to the underlying disease mechanism. **(B)** Summary table of the known actin and microtubule effects of profilin-1 (wild-type), each of the ALS-associated profilin variants, and the actin-binding (i.e., R88E and H120E) and poly-*L*-proline (Y6D) binding-deficient controls.

Wild-type profilin has efficient folding dynamics and can refold following several denaturization procedures (Kaiser et al., 1989). Several studies have explicitly detailed folding and stability defects with certain ALS-linked profilin variants (Boopathy et al., 2015; Del Poggetto et al., 2016; Freischmidt et al., 2015; Kiaei et al., 2018; Pereira et al., 2019; Sadr et al., 2021). The solubility and stability of each profilin variant is an important consideration for this study. Some ALS-linked mutations, particularly C71G, are destabilizing (Figure 3) (Boopathy et al., 2015; Figley et al., 2014; Lim et al., 2016). Are these folding issues part of the mechanism of ALS or merely the result of lower effective concentrations or less active profilins? The A20T, M114T, G118V, and R136W variants each exhibited gain of function effects in some of the assays used to test actin activities above. C71G appears to block actin filament nucleation closer to wild-type levels. Q139L may be more effective at blocking nucleation, though the observed difference is not statistically significant. Protein instability or differences in experimental design may explain inconsistencies regarding the role of C71G in mDia1-related actin assembly (Figure 4), although in different experiments this protein was able to bind PLP-peptide (Figure 5) (Schmidt et al., 2021). In contrast, C71G has been found in cellular protein degradation pathways which may support the notion that misfolding contributes to this variant’s pathological mechanism (Figley et al., 2014; Pohl and Dikic, 2019).

Profilin regulates actin in two opposing ways. First, it binds to actin monomers and effectively suppresses actin filament nucleation (Davey and Moens, 2020; Pimm et al., 2020; Skruber et al., 2018). Second, it can stimulate actin assembly through filament nucleation and/or elongation mechanisms where profilin simultaneously binds monomers and the PLP regions of other proteins like formins or ENA/VASP (Ferron et al., 2007; Funk et al., 2019; Skruber et al., 2020; Suarez and Kovar, 2016). Our results demonstrate that each of the ALS-variants influences the affinity of profilin for actin monomers (Figure 1C). These results likely explain the increase in actin filament nucleation of several ALS-variants compared to wild-type profilin observed in TIRF assays. Each variant was still able to significantly suppress actin nucleation compared to controls containing polymerizing actin filaments alone. We did not observe any changes to actin filament elongation rates, with the exception of a subtle change to A20T that might be explained with changes to actin alone controls between experiments. The A20 residue is located opposite to the actin binding surface on profilin yet binds actin monomers weaker than wild-type. It was also more prone to chemical denaturation, thus subtle destabilization induced by this mutation may explain these effects.

The inhibition of actin assembly by profilin is a well characterized facet of profilin-based actin regulation in vitro. However, lipid-based or formin-mediated actin assembly mechanisms stimulated by profilin dominate cellular actin dynamics to maintain neuronal homeostasis, cellular morphologies, and signal transduction (Figure 6A)(Pimm et al., 2020; Pinto-Costa et al., 2020; Rotty et al., 2015; Skruber et al., 2018; Suarez et al., 2015; Zimmermann et al., 2017). Here we demonstrate that half of the identified ALS-associated profilin variants are impaired in formin-mediated actin assembly. Each of the ALS-profilins bind a small peptide of mDia1 containing two PLP motifs with similar affinities (Figure 5B). Several variants (A20T, T109M, R136W, and Q139L) are located on or near the surface of profilin that interacts with PLP. Most of these variants lack the ability to stimulate formin-based actin assembly despite binding a small mDia1-derived PLP peptide with similar affinity to the wild-type protein. It is difficult to discern if this is biologically relevant or a confounding effect due to the presence of only two of the fifteen mDia1 binding sites. Opposite to these effects, the M114T and G118V variants also bound PLP peptides with slightly stronger affinity than wild-type profilin and somehow enhance actin elongation by mDia1 (Figures 4C and 5B). Stimulation of formin-based actin filament polymerization by the G118V variant has similarly been noted by others (Schmidt et al., 2021). The combination of strong PLP binding and weakened actin association may serve to enhance polymerization by decreasing the dwell time of the profilin-actin complex upon addition to a growing filament (Zweifel et al., 2021).

The mechanism used by formins to accelerate actin polymerization is complex and not fully elucidated. In general terms, the conformation of formin homology 2 domains encircling the growing actin filament contributes to the overall polymerization speed, with those having a more open, looser conformation tending to polymerize faster (Aydin et al., 2018; Breitsprecher and Goode, 2013; Courtemanche, 2018; Henty-Ridilla et al., 2016). PLP-containing FH1 “arms” use profilin-bound actin monomers to expedite assembly, likely by increasing the local actin concentration and/or by orienting actin monomers for efficient incorporation into the growing filament (Breitsprecher and Goode, 2013; Cao et al., 2018; Chesarone et al., 2010; Courtemanche and Pollard, 2012; Funk et al., 2019; Homa et al., 2021; Zweifel and Courtemanche, 2020). The number and location of PLP tracts in the FH1 domain also contribute to this mechanism (Courtemanche, 2018; Paul et al., 2008; Zweifel and Courtemanche, 2020). Elegant studies examining the yeast mDia1 homolog demonstrate that competitive interactions along the PLP tracts ultimately deliver profilin-bound actin to the site of assembly and must also efficiently unbind to clear the way for the next building block (Courtemanche, 2018; Zweifel and Courtemanche, 2020). It is currently unclear which of these mechanisms (or others) could explain the effects of the ALS-related mutations in profilin. In addition, there are fifteen mammalian formins, Ena/VASP, and additional PLP-containing ligands known to use profilin to gain access to cellular actin pools. Toward this end, compared to wild-type profilin the C71G, M114T, and G118V variants seem to favor binding specific formins mDia1, mDia2, and FMNL1 in cells (Schmidt et al., 2021). Exactly how cellular actin and profilin pools are regulated and what proportion is available for diverse functions or interactions remain intriguing open questions.

What does the observation that certain profilin variants prefer specific formins mean for profilin in neurons or neurodegenerative disease states? In addition to its role with the actin cytoskeleton or with various cellular lipids, profilin also directly binds and regulates the dynamics of tubulin and microtubules (Henty-Ridilla et al., 2017; Pimm et al., 2021). Although C71G has not been tested, both M114T and G118V lack for microtubule-based activities (Henty-Ridilla et al., 2017). Thus, one intriguing possibility is that these ALS-variants display higher concentrations bound to formins because they are liberated from microtubules (Henty-Ridilla et al., 2017; Schmidt et al., 2021). A study linking the M114T and G118V variants to formins did not detect tubulin or microtubules from immunoprecipitation experiments with wild-type profilin or the ALS-associated variants (Schmidt et al., 2021). This is surprising because genetic and biochemical evidence strongly suggest profilin-microtubule interactions, but might be explained from temperature-dependent effects, lysis conditions or sensitivity of profilin to N-terminal tags which can influence binding interactions (Figley et al., 2014; Nejedla et al., 2017, 2016; Nejedlá et al., 2021; Pimm et al., 2021; Schmidt et al., 2021; Witke et al., 1998; Wittenmayer et al., 2004). A newly developed tool permits the visualization of a subset of cellular profilin molecules (Pimm et al., 2021). Consistent with microtubule binding affecting cellular interactions, we have observed that G118V mutant profilin shows reduced association with microtubules in cells, while a mutant that blocks formin interaction shows enhanced association (Pimm et al., 2021). This tool may be helpful in future cellular experiments of these complicated and exciting profilin variants or additional studies linking profilin to other forms of neurodegeneration.

## Acknowledgements

We are grateful to Marc Ridilla (Repair Biotechnologies) for helpful comments on this manuscript. We also appreciate past discussions with Bruce L. Goode (Brandeis University), Daryl Bosco (University of Massachusetts Medical School), John Landers (University of Massachusetts Medical School), and Sivakumar Boopathy (Harvard Medical School). This work was supported by the National Institutes of Health [GM133485]; ALS Association, Washington, DC [starter grant 20-IIP-506] and a scholar award from the Alexandrine and Alexander Sinsheimer Fund, Chicago, IL.

## Author contributions

XL, AB, BH, MLP, JLH-R purified proteins. BH, JLH-R designed experiments. XL, AB, MLP, JLH-R performed analyses. JLH-R made the figures. XL, BH, MLP, JLH-R wrote and edited the paper.

## Supplemental Information

**Figure S1.**
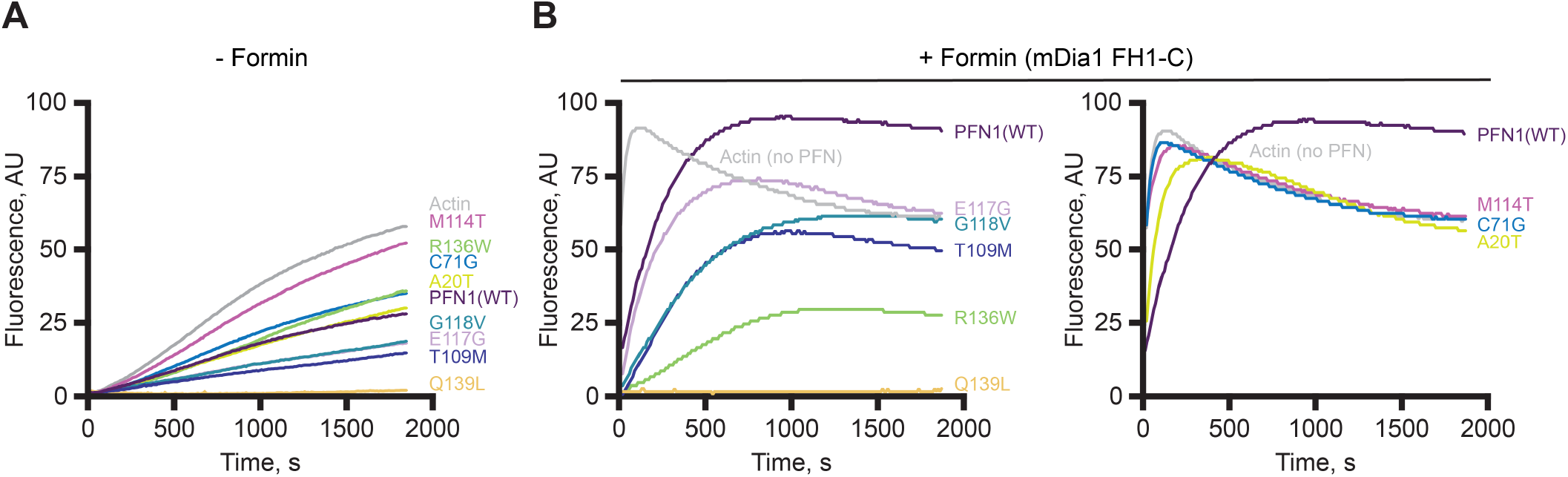
ALS-associated profilin variants affect bulk actin filament assembly. **(A)** Mean fluorescence intensity traces from pyrene fluorescence assays containing 2 μM actin (5% pyrene-labeled) polymerized in the presence of 5 μM profilin (wild-type (PFN1) or ALS-variant). Traces are the average intensity over time for three separate experiments. **(B)** Reactions as in (A) supplemented with 25 nM constitutively active formin, mDia1(FH1-C). Separated traces in (B) are for clarity. Left panel contains variants with weakened formin-mediated actin polymerizing activity; Right panel contains ALS-variants that do not stimulate formin-polymerization. The actin and wild-type profilin curves are duplicated in the left and right panels for ease of comparison. Values in (A) and (B) were performed on the same plate per replicate.

**Figure S2.**
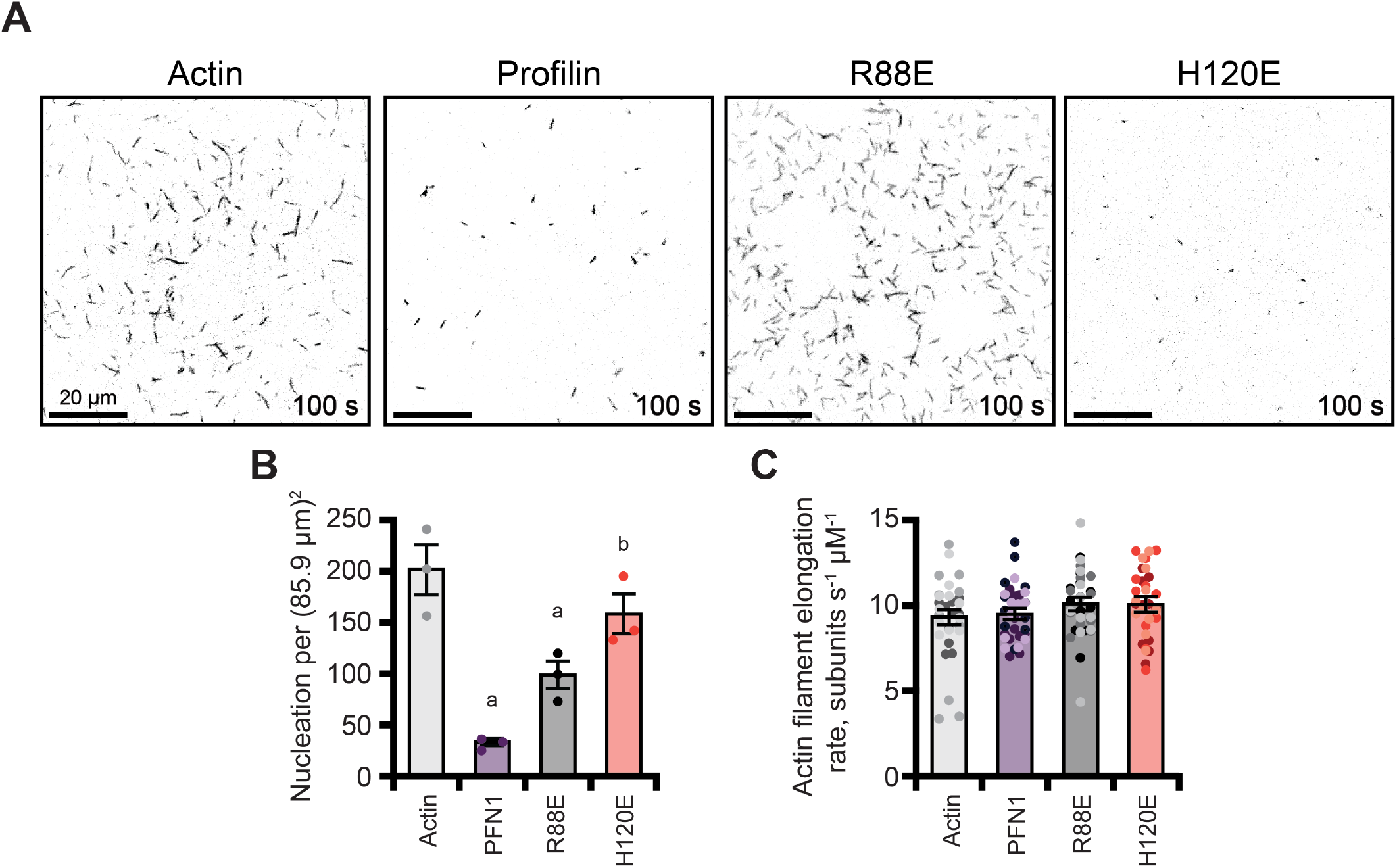
Actin binding deficient profilin proteins do not efficiently suppress actin filament nucleation. **(A)** Representative fields of view (FOV) from TIRF microscopy assays containing 1 μM actin (10% Oregon Green (OG)-labeled, 0.6 nM biotin-actin) assembled in the absence (actin) or presence of 5 μM profilin (wild-type (PFN1), R88E, or H120E). Scale bars, 20 μm. **(B)** Nucleation measurement (mean count) of actin filaments visible from TIRF reactions shown in (A). Measurements were made 100 s after the actin polymerization was initiated. Each dot represents counts from different experimental replicates. **(C)** Distribution of actin filament elongation rates from TIRF reactions as in (A). Dots represent individual measurements (n = 11 per replicate or n = 33 total). Shaded values show the distribution of different independent experimental replicates (n = 3). Error bars indicate SEM. a, significantly different (P < 0.05) than actin control; b, significantly different (P < 0.05) than wild-type profilin control. Number of measurements determined by power analysis. Significant differences for nucleation experiments were determined by one-way ANOVA with Bonferroni post-hoc analysis. For elongation rate experiments, Tukey post-hoc analysis was used.

**Figure S3.**
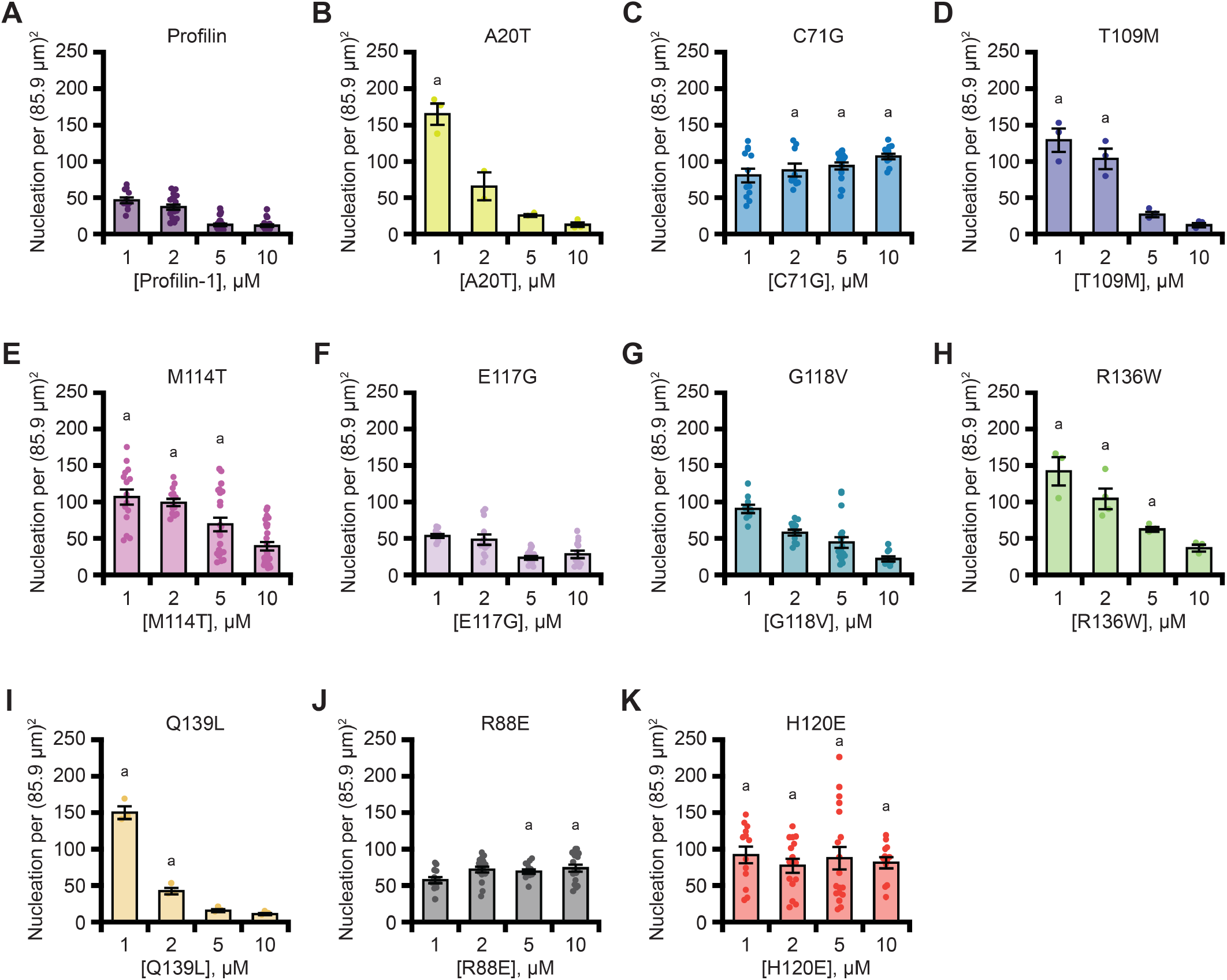
Dose dependent inhibition of actin filament nucleation by profilin (wild-type and ALS-relevant variants). Nucleation measurement (mean count) of actin filaments visible from TIRF reactions containing 1 μM actin (10% Oregon Green (OG)-labeled, 0.6 nM biotin-actin) assembled in the presence of either 1 μM, 2 μM, 5 μM, or 10 μM profilin (wild-type or mutants). Measurements were made 240 s after the actin polymerization was initiated. Each dot represents counts from different experimental replicates. Legend is as follows: **(A)** Wild-type profilin **(B)** A20T **(C)** C71G **(D)** T109M **(E)** M114T **(F)** E117G **(G)** G118V **(H)** R136W **(I)** Q139L. The R88E **(J)** and H120E **(K)** mutants are negative controls (not ALS-associated variants) that do not bind actin well. Error bars indicate SEM. a, significantly different (P < 0.05) than wild-type profilin of the same concentration. Significant differences determined by one-way ANOVA with Bonferroni post-hoc analysis.

**Figure S4.**
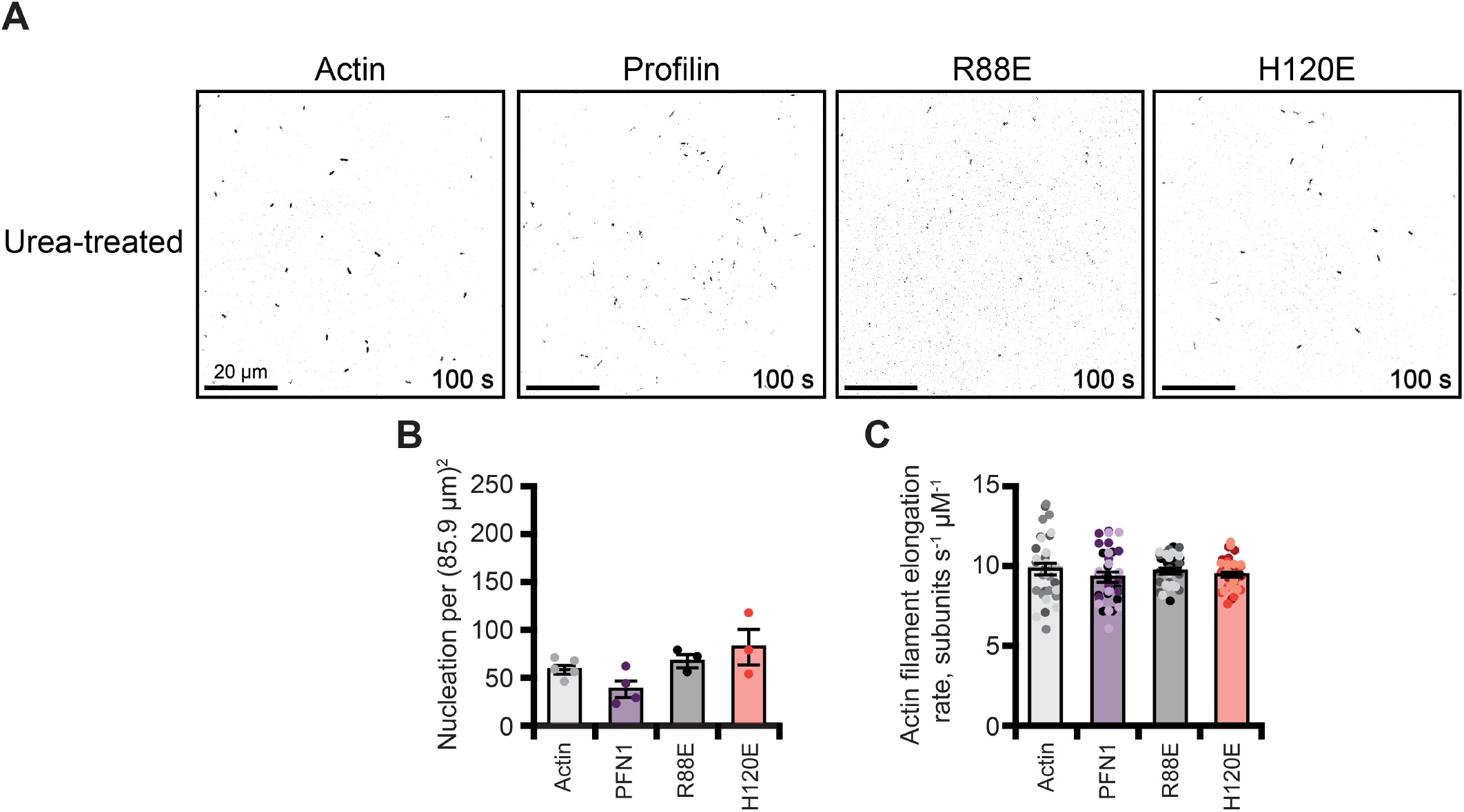
Effects of urea denaturation on actin-binding impaired profilin mutants. **(A)** Representative fields of view (FOV) from TIRF microscopy assays containing 1 μM actin (10% Alexa-488 labeled, 0.6 nM biotin-actin) in the absence (control) or presence of 5 μM profilin (wild-type, R88E, or H120E) that was treated with 5 M urea for 30 min, diluted to 344 mM in TIRF buffer, and allowed to refold for 30 min. After the refolding period urea-treated profilins were assessed for actin activities in TIRF assays. The final concentration of urea in each of these experiments (including actin controls) was 344 mM. Scale bars, 20 μm. **(B)** Mean count of actin filaments 100 s after actin polymerization was initiated as in (A). Each dot represents counts from independent experimental replicates. **(C)** Distribution of actin filament elongation rates from TIRF reactions as in (A). Dots represent individual measurements (n = 11 per replicate or n = 33 total). Shaded values show the distribution of different independent experimental replicates (n = 3). Error bars indicate SEM. a, significantly different (P < 0.05) than actin control; b, significantly different (P < 0.05) than wild-type profilin control. Number of measurements determined by power analysis. Significant differences for nucleation experiments were determined by one-way ANOVA with Bonferroni post-hoc analysis. For elongation rate experiments, Tukey post-hoc analysis was used.

**Figure S5.**
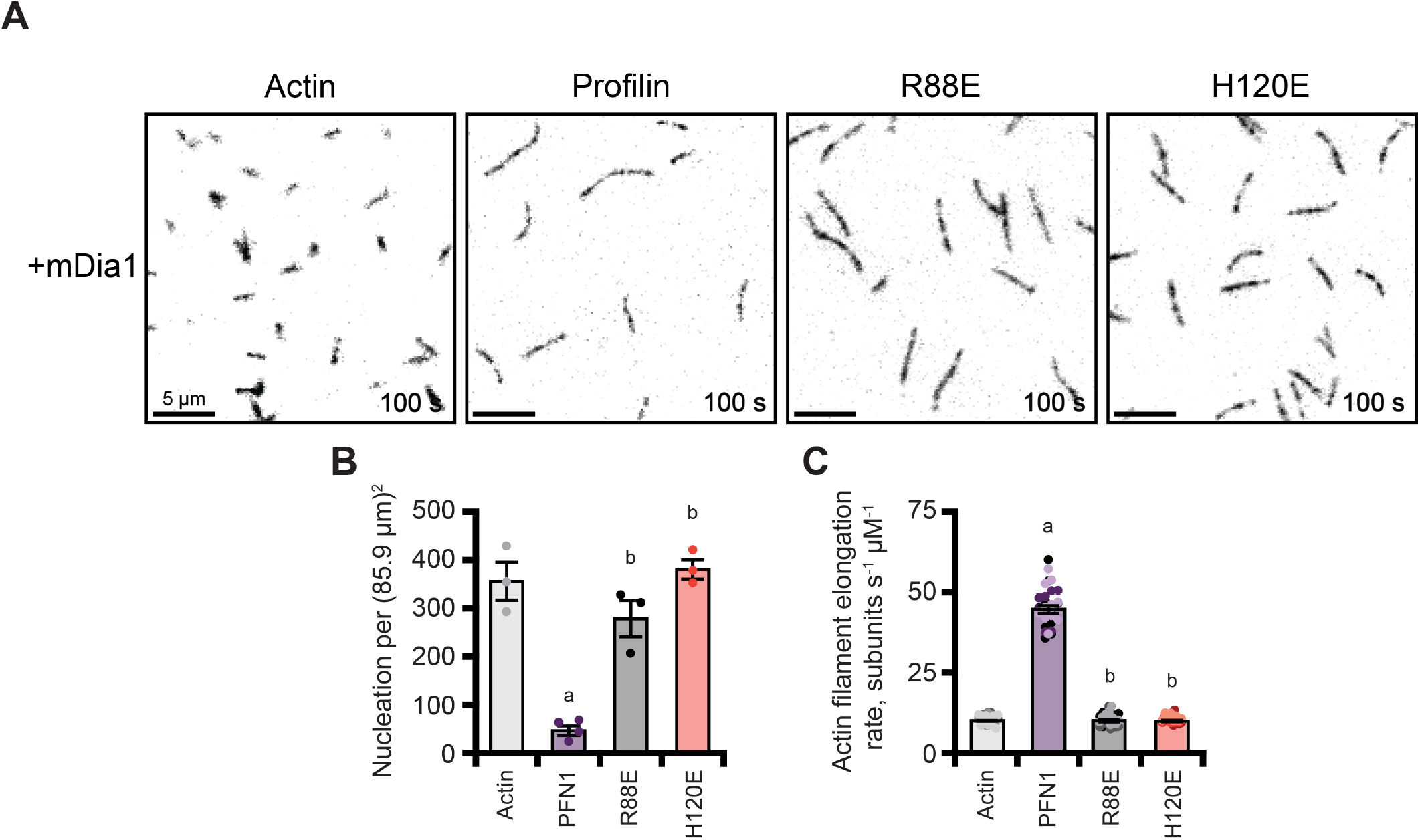
Actin-binding impaired profilin mutants do not stimulate formin-based actin polymerization. **(A)** Representative time lapse TIRF images from actin polymerization assays in the presence of formin and absence or presence of profilin. Reactions contain 1 μM actin (10% Alexa-488 labeled, 0.6 nM biotin-actin) and 10 nM mDia1(FH1-C) without or with 5 μM profilin (wild-type, R88E, or H120E). **(B)** Graphical representation of actin filament nucleation for TIRF movies as in (A); individual data points represent the number of filaments per field of view at 100 s after initiation of actin assembly from separate polymerization experiments. Error bars, represent SEM. **(C)** Distribution of actin filament elongation rates from TIRF reactions as in (A). Dots represent individual measurements (n = 11 per replicate or n = 33 total). Shaded values show the distribution of different independent experimental replicates (n = 3). Error bars indicate SEM. a, significantly different (P < 0.05) than actin control; b, significantly different (P < 0.05) than wild-type profilin control. Number of measurements determined by power analysis. Significant differences for nucleation experiments were determined by one-way ANOVA with Bonferroni post-hoc analysis. For elongation rate experiments, Tukey post-hoc analysis was used.

## Supplemental movies

**Supplemental Movie 1. TIRF microscopy comparing the effects of each ALS-associated profilin protein on actin assembly**. Images were acquired at 5 s intervals. Reaction components: 1 μM actin monomers (10% Oregon Green (OG)-labeled; 0.6 nM biotin-actin). Variable components 5 μM profilin (wildtype or ALS-variant). Video playback is 10 frames per s. Scale bars, 10 μm.

**Supplemental Movie 2. TIRF microscopy comparing the effects of each ALS-associated profilin on actin assembly following treatment with 5 M urea**. Images were acquired at 5 s intervals. Reaction components: 1 μM actin monomers (10% Oregon Green (OG)-labeled; 0.6 nM biotin-actin). Variable components 5 μM profilin (wildtype or ALS-variant). Video playback is 10 frames per s. Scale bars, 10 μm.

**Supplemental Movie 3. TIRF microscopy comparing the effects of the ALS-associated profilins with formin**. Images were acquired at 5 s intervals. Reaction components: 1 μM actin monomers (10% Alexa-488 labeled; 0.6 nM biotin-actin) and 10 nM mDia1(FH1-C). Variable components 5 μM profilin (wildtype or ALS-variant). Video playback is 10 frames per s. Scale bars, 10 μm.

